# Attention recruits frontal cortex in human infants

**DOI:** 10.1101/2020.10.14.340216

**Authors:** C. T. Ellis, L. J. Skalaban, T. S. Yates, N. B. Turk-Browne

**Affiliations:** Department of Psychology, Yale University, New Haven, CT 06511

## Abstract

Young infants learn about the world by overtly shifting their attention to perceptually salient events. In adults, attention recruits several brain regions spanning the frontal and parietal lobes. However, these regions are thought to have a protracted maturation and so it is unclear whether they are recruited in infancy and, more generally, how infant attention is supported by the brain. We used event-related fMRI with 24 awake behaving infants 3–12 months old while they performed a child-friendly attentional cuing task. A target was presented to either the left or right of the infant’s fixation and eye-tracking was used to measure the latency with which they saccaded to the target. To manipulate attention, a brief cue was presented before the target in three conditions: on the same side as the upcoming target (valid), on the other side (invalid), or on both sides (neutral). All infants were faster to look at the target on valid versus invalid trials, with valid faster than neutral and invalid slower than neutral, indicating that the cues effectively captured attention. We then compared the fMRI activity evoked by these trial types. Regions of adult attention networks activated more strongly for invalid than valid trials, particularly frontal regions such as anterior cingulate cortex. Neither behavioral nor neural effects varied by infant age within the first year, suggesting that these regions may function early in development to support the reorienting of attention. Together, this furthers our mechanistic understanding of how the infant brain controls the allocation of attention.

---

Having an attention system that is capable of swiftly reorienting to salient events (i.e., stimulus-driven attention) is essential for many behaviors. This is perhaps most true in infancy, during which exploration is thought to be critical (Gibson, 1988) and attention allows infants to fully experience learning moments (Baillargeon, 1987). The value of attention in early development might explain why infants are equipped with the capacity to flexibly allocate attention: They can saccade to onsets soon after birth (Aslin and Salapatek, 1975), use cues to facilitate orienting (Farroni et al., 2004; Johnson et al., 1994), and make predictions about upcoming events (Emberson et al., 2015). Yet, how the infant brain supports attention remains a mystery.

An extensive literature in adults could inform our understanding of the neural basis of stimulus-driven attention in infants. In adults, attention is supported by the ventral and dorsal frontoparietal networks, consisting of the right temporal parietal junction (TPJ), superior parietal lobe (SPL), lateral occipital cortex (LOC), frontal eye fields (FEF), and middle/inferior frontal gyrus (MFG/IFG) (Arrington et al., 2000; Corbetta et al., 2008; Corbetta and Shulman, 2002; Doricchi et al., 2010), and the cingulo-opercular network, consisting of the anterior cingulate cortex (ACC), insula, and basal ganglia (Dosenbach et al., 2007; Shulman et al., 2009). However, these regions are anatomically immature in infants (Casey et al., 2005; Gilmore et al., 2012; Gogtay et al., 2004; Matsuzawa et al., 2001) and functional connectivity between these regions, critical for supporting attention in adults (Farrant and Uddin, 2015; Konrad et al., 2005), is weak in early infancy (Gao et al., 2015). For example, the connectivity of ACC emerges over early development ([Gao et al. 2015, 2009] but see [Eyre et al. 2020]). Indeed, these regions undergo functional changes late into adolescence (Farrant and Uddin, 2015; Konrad et al., 2005). Furthermore, the adult frontoparietal network is also recruited for maintaining goals and volitionally directing attention based on them (i.e., goal-directed attention) (de Fockert et al., 2004; Kincade et al., 2005; Meyer et al., 2018). However, goal-directed attention is less developed than stimulus-driven attention in infants (Csibra et al., 1998; Hood et al., 1998; Jakobsen et al., 2013; Scaife and Bruner, 1975), further suggesting immaturity in some parts of infant attention networks. These threads of evidence led to the proposal that some regions, like the ACC, are not sufficiently mature in infancy to support attention, and instead the TPJ, SPL and FEF are recruited (Posner et al., 2012). An alternative account suggests that the frontoparietal and cingulo-opercular networks are capable of functioning in early infancy (Eyre et al., 2020), even if there is anatomical immaturity (Raz and Saxe, 2020).

Existing studies of the infant attention system have been inconclusive about the extent to which the infant brain recruits adult-like attention networks. Electroencephalography (EEG) with infants suggested that some neural signatures of attention are adult-like (Xie et al., 2018; Xie and Richards, 2017). However, EEG has insufficient spatial resolution to resolve which regions are supporting attention. Functional near-infrared spectroscopy (fNIRS) offers potentially greater resolution (Emberson et al., 2015; Lloyd-Fox et al., 2019), but is unable to localize activity beyond regions close to the scalp surface, including deeper, ventral, medial, and subcortical structures such as the ACC and basal ganglia. Hence, it remains unclear how stimulus-driven attention is supported in the infant brain.

We used fMRI and eye-tracking to investigate the behavioral and neural basis of stimulus-driven attention in awake behaving infants under a year old. Among non-invasive techniques, fMRI is uniquely capable of resolving brain-wide, fine-grained attention processes. Recent innovations have made it possible to collect data from awake behaving infants (Biagi et al., 2015; Deen et al., 2017; Dehaene-Lambertz et al., 2002; Ellis et al., 2020b). Using a developmental variant (Johnson et al., 1994) of the Posner cuing task (Posner et al., 1978), we simultaneously recorded gaze behavior and whole-brain fMRI activity to uncover the neural basis of attention in infants.

## Results

We administered a cuing task while simultaneously collecting fast event-related fMRI to examine how the brain supports stimulus-driven attention in 24 awake behaving infants aged 3–12 months old (Table S1). On each trial, participants saw a brief cue followed by a salient, spinning pinwheel target (Johnson et al., 1994). To assess behavioral evidence of attention, we measured the response time (RT) of saccades to the target (Figure 1A). This target appeared on the left or right side of the display equally often. On 50% of trials the target was preceded by a cue that appeared on the same side as the target (valid trials). On 25% of trials the cue appeared on the opposite side as the target (invalid trials) and on the remaining 25% of trials the cue appeared on both sides (neutral trials).

**Figure 1:**
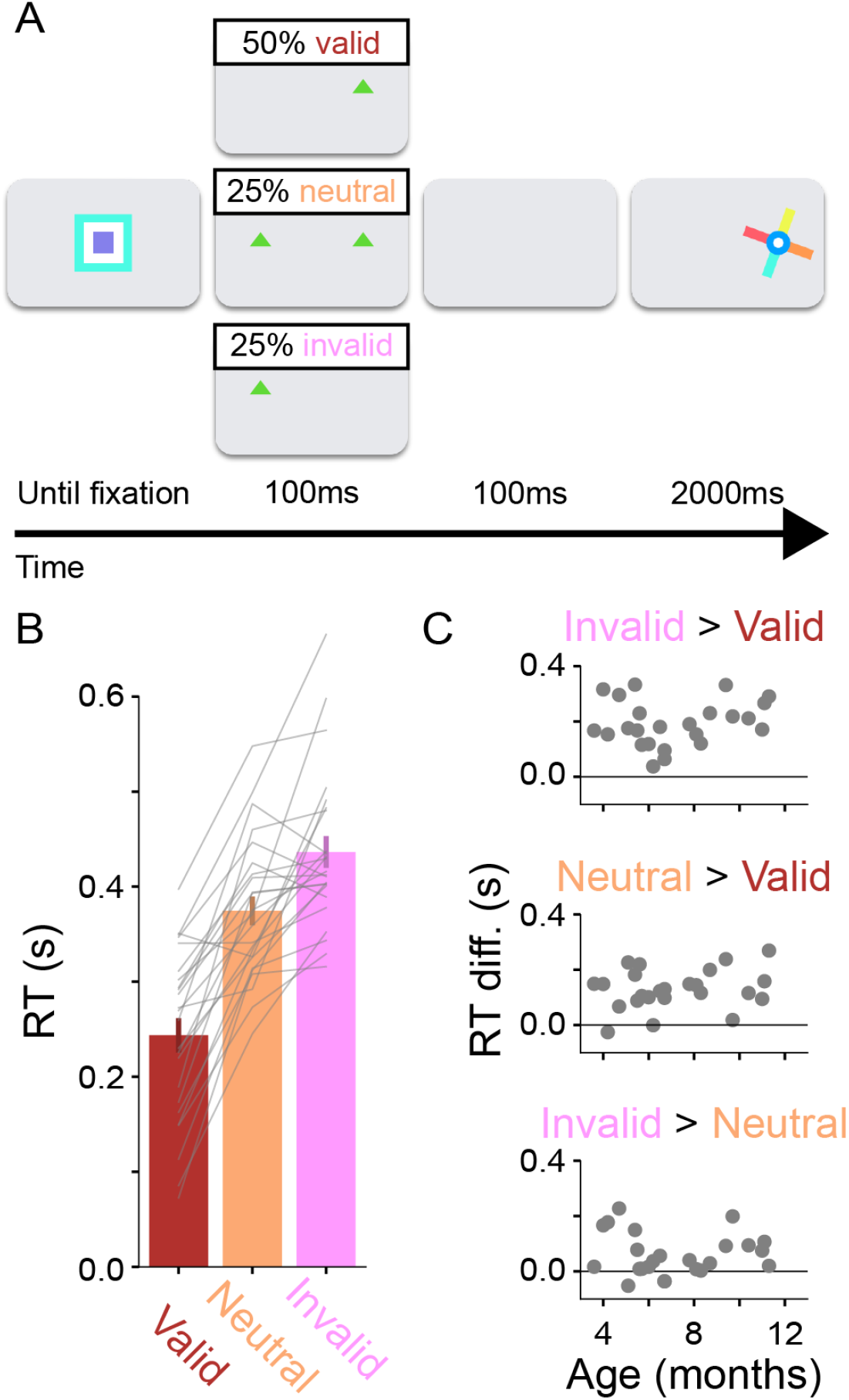
Task design and behavioral evidence of stimulus-driven attention in infants. (A) Trial sequence: Participants were presented with an attention-getter until they fixated. A cue was then presented briefly before a target appeared. The cue was either in the same location as the target (valid), in the opposite location (invalid), or cues were presented bilaterally (neutral). (B) Average RT in seconds for each trial type. Error bars indicate standard error across participants. Grey lines connect individual participant data across the three trial types. (C) Relationship of different condition comparisons to age in months.

We quantified the RT to the target for each of these trial types separately. Our main behavioral index of stimulus-driven attention was the RT difference between invalid and valid trials (Figure 1B). All participants looked to the target faster on average on valid than invalid trials (invalid>valid in 24/24 participants, *M*=0.19s, 95% CI=[0.16, 0.22], *p*<0.001). This difference could be driven by valid cues facilitating attention, invalid cues inhibiting attention, or both. We found evidence for facilitation by comparing valid with neutral trials (neutral>valid in 23/24 participants, *M*=0.13s, CI=[0.10, 0.16], *p*<0.001). We also found evidence for inhibition by comparing invalid with neutral trials (invalid>neutral in 21/24 participants, *M*=0.06s, CI=[0.03, 0.09], *p*<0.001). These effects were seen even in the first block of the task (Figure S1), suggesting that this behavior did not depend on learning that the cues predicted the target location (because of the higher probability of valid than invalid or neutral).

An important consideration is the speed of the saccades, especially on valid trials. It typically takes 300–500ms for an infant to initiate and complete a saccade (Aslin and Salapatek, 1975; Johnson et al., 1994), yet a large proportion of trials had RTs much faster than that (Figure S2A). This suggests that infants initiated a saccade immediately after the cue appeared, rather than wait for the target to appear. Nevertheless, what is important is that these cues were robust drivers of stimulus-driven attention, showing that infants are spontaneously utilizing the cues to allocate attention.

The attention effects observed in the full sample did not vary by age (Figure 1C). Namely, the RT differences between trial types were not reliably correlated with age in months (invalid>valid: r=0.17, *p*=0.352; neutral>valid: r=0.23, *p*=0.314; invalid>neutral: r=−0.03, *p*=0.873). That said, older participants were faster overall (Figure S2B; r=−0.63, *p*<0.001), demonstrating that we had sensitivity in principle to detect large age effects. We are cautious about drawing a definitive conclusion from null effects, and a larger sample size may reveal a more subtle relationship. However, these findings are consistent with the possibility that younger and older infants show similar cuing effects.

We investigated the neural mechanisms supporting stimulus-driven attention in infancy using a univariate general linear model (GLM) that contrasted evoked responses for the valid, neutral, and invalid trials. The premise of this analysis is that brain regions that contribute to attentional processing will be activated when attention is recruited. The behavioral results show that the cue oriented attention on both valid and invalid trials (relative to neutral). However, the longer RTs on invalid trials suggest additional attentional processes were needed to support reorienting. The neutral trials also provide an interesting baseline, as attention may not have been allocated until after the appearance of the target. Hence contrasts between these trial types can reveal different mechanisms supporting stimulus-driven attention (Doricchi et al., 2010).

Our primary analyses involve performing these contrasts within regions of interest (ROIs) from adult attention networks. However, given the challenge of awake infant fMRI, as a first pass we conducted a voxelwise whole-brain analysis to verify that we were able to collect data of reasonable quality and that the cuing task drove responses in the infant brain. For each comparison between trial types, we calculated the *t*-statistic across participants in each voxel (Figure 2). Widespread activity that distinguished between trial types was observed at a liberal threshold, which we next quantify with greater precision and sensitivity by averaging across voxels within key ROIs.

**Figure 2:**
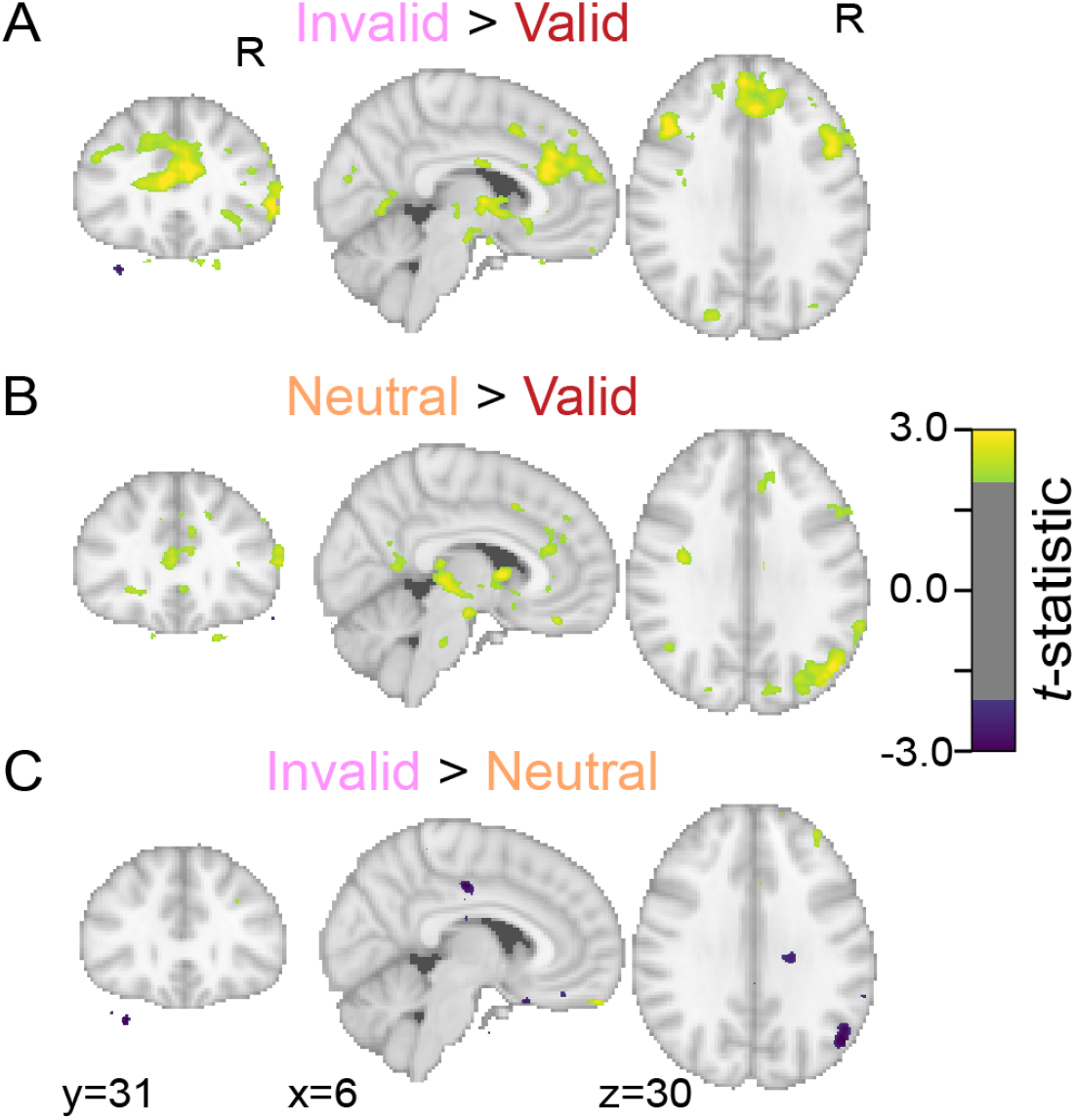
Whole-brain statistical map of difference between conditions. Contrasts for each participant were aligned to standard space and tested for reliability with a group *t*-test. Three tests were performed: (A) invalid>valid, (B) neutral>valid, and (C) invalid>neutral. An uncorrected threshold is used for visualization (*p*<0.05). Coordinates are in adult MNI space.

To test whether regions from adult attention networks were recruited in infants, we defined seven ROIs (Table S2) independently based on a functional atlas (Yarkoni et al., 2011) and compared each contrast across participants (Figure 3). The contrast of valid and invalid trials, which tests an overall effect of attentional cuing, resulted in significantly higher activity for invalid trials in LOC (*M*=0.47, CI=[0.05, 0.87], *p*=0.032), right ACC (*M*=0.37, CI=[0.04, 0.70], *p*=0.027), and right MFG (*M*=0.40, CI=[0.05, 0.76], *p*=0.028), and marginally higher activity in right TPJ (*M*=0.40, CI=[-0.03, 0.79], *p*=0.066). The contrast of valid and neutral trials, which tests the facilitatory effects of a cue, there was significantly higher activity for neutral trials in right ACC (*M*=0.37, CI=[0.03, 0.69], *p*=0.033), and LOC (*M*=0.47, CI=[0.09, 0.84], *p*=0.017), and marginally higher activity in IPS (*M*=0.38, CI=[0.00, 0.72], *p*=0.051). The contrast of invalid and neutral trials, which tests the inhibitory effects of a cue, resulted in no significant regions. In sum, regions thought to support attention in infancy, such as SPL and FEF (Posner et al., 2012), were not recruited for stimulus-driven attention, whereas regions thought to be immature, did support stimulus-driven attention, particularly the facilitation of processing by cues.

**Figure 3:**
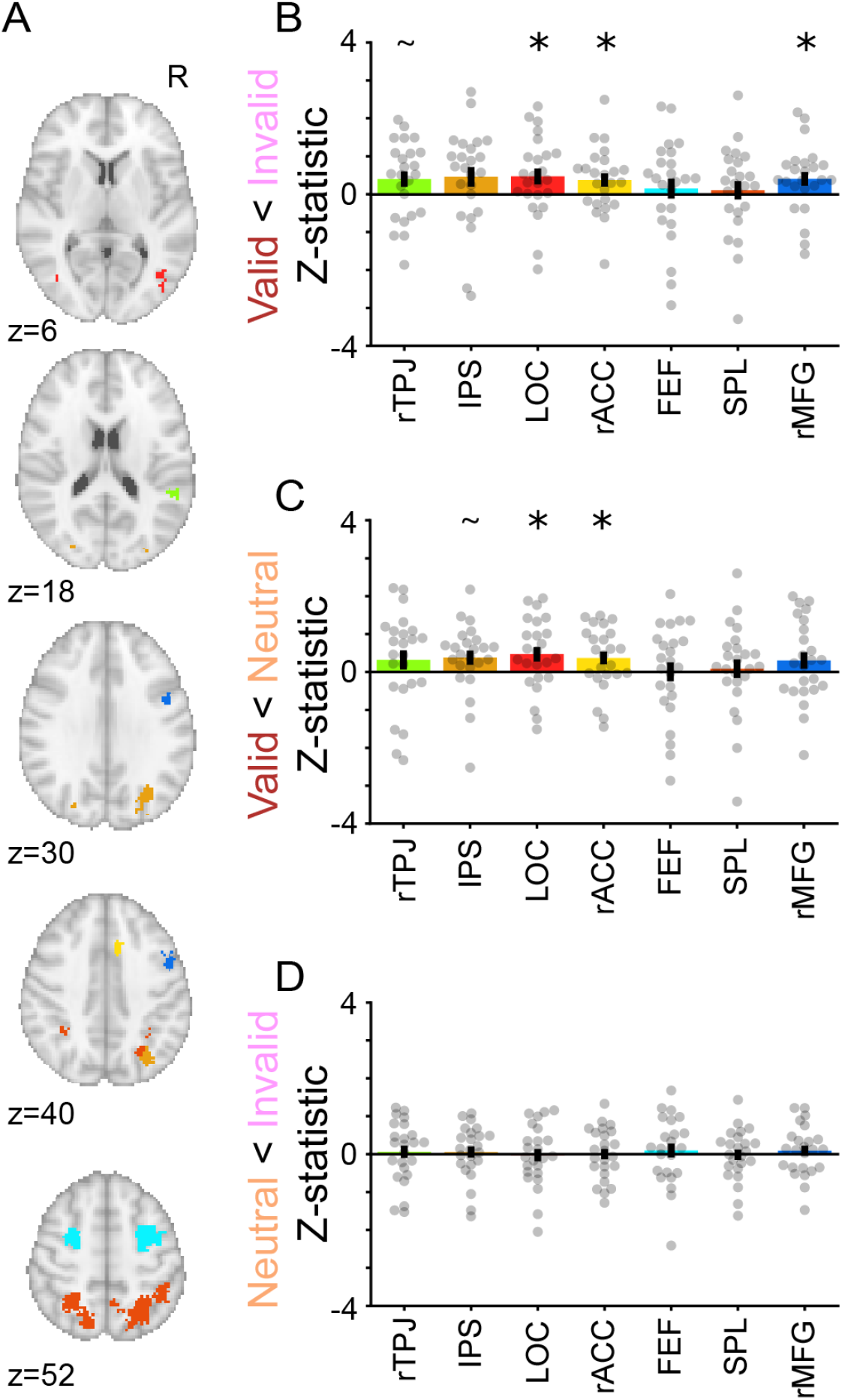
Neural evidence of stimulus-driven attention in infants. (A) Individual ROIs from functional analysis with coordinates in MNI space. Contrasts were extracted from these ROIs, averaged across voxels, and tested for reliability with bootstrap resampling. Three tests were performed: (B) invalid>valid, (C) neutral>valid, and (D) invalid>neutral. Lower-case ‘r’ and ‘l’ indicate left and right hemisphere, respectively. Error bars indicate standard error across participants. *=*p*<0.05, ^~^=*p*<0.075

Some of these ROIs were small, which could lead to unstable estimates of the average evoked response in different conditions and explain why only a subset of the regions reached significance in different contrasts. However, dilating the ROIs to increase their size yielded very similar results (Figure S3). Moreover, if we use a spherical ROI around the peak of each functional ROI, thus balancing the number of voxels averaged per region, the results are also similar (Figure S4). Together, these findings suggest that frontal, occipital, and, to a lesser extent, parietal regions are recruited to support stimulus-driven attention in infants. We did not observe any evidence that the strength of the invalid>valid effect correlated with participant age (Figure S5).

The analyses above consist of ROIs defined from meta-analyses of adult fMRI studies, and thus assume adult anatomy and that there is a consistent mapping from adults to infants in MNI standard space. To avoid such assumptions, we performed an additional analysis using cross-validation within the infants to define ROIs in an unbiased, data-driven way. Specifically, we performed a leave-one-participant-out analysis in which clusters of voxels were defined in all but one participant and then the evoked response from those clusters were quantified in the remaining participant (Figure S6; Esterman et al. 2010). Even though we did not make assumptions about where we would find differences, these analyses implicated similar regions in the frontal lobe, including the anterior cingulate cortex and inferior frontal gyrus. Specific to this analysis, we found involvement of the basal ganglia, especially the caudate and thalamus, which is part of the cingulo-opercular attention network (Dosenbach et al., 2007).

In this task, infants allocated attention with eye movements, so it is possible that the differences between conditions were driven by eye movements rather than attention *per se* (Perry and Zeki, 2000). In particular, more saccades were made on invalid trials (*M*=2.10) compared with valid (*M*=1.75, difference CI=[0.20, 0.50], *p*<0.001) and neutral trials (*M*=1.84, difference CI=[0.10, 0.42], *p*=0.001). To evaluate this possibility, we tested whether saccades themselves evoked responses in the ROIs defined previously (Figure 3A). We ran a separate GLM with a saccade regressor in which events were defined every time the participant moved their eyes. None of the attention ROIs were significantly activated by saccades (Figure S7B). As a positive control to validate this analysis, non-attention regions in early visual cortex were activated (Figure S7A; Tse et al. 2010).

## Discussion

We investigated the neural mechanisms of stimulus-driven attention in infants under the age of 12 months. We found robust behavioral evidence that infants allocated attention to cues in our task. We tested whether this attention behavior was supported by regions of frontoparietal and cingulo-opercular networks that are involved in analogous tasks in adults. Posing a challenge to existing theories (Posner et al., 2012), infant attention recruited regions of the cingulo-opercular network, such as the ACC and basal ganglia, as well as anterior portions of the frontoparietal network, including the MFG/IFG. This suggests that the human brain may have a surprisingly functional frontal cortex in the first year of life.

The engagement of regions from well-characterized attention networks may help interpret our findings, based on the functions of these regions in adults. In the frontal lobe, the MFG/IFG (Corbetta and Shulman, 2002; de Fockert et al., 2004; Doricchi et al., 2010; Kincade et al., 2005; Shulman et al., 2009) and the ACC (Dosenbach et al., 2007; Shulman et al., 2009) are recruited for reorienting attention. These regions are part of the ventral frontoparietal network (Corbetta and Shulman, 2002) or salience network (Touroutoglou et al., 2012). The MFG/IFG may serve as a highly-connected node for distributing information in the frontoparietal network (Fox et al., 2006). The involvement of the ACC is more surprising, given its role in cognitive control (Miller and Cohen, 2001) and goal maintenance (Dosenbach et al., 2007) in adults. These functions have protracted developmental trajectories into late adolescence (Munakata et al., 2012). Indeed, the ACC is only weakly connected at rest to other parts of the brain in the first year of life (Gao et al., 2015, 2009; Posner et al., 2012). This raises the interesting possibility that the function of the ACC may be different in infants compared with adults. Evidence of the recruitment of these regions was stable across the age range, though future studies with larger sample sizes would be needed if the age relationship was small or moderate in size. Regardless, despite the protracted anatomical development of the frontal lobe (Casey et al., 2005; Gilmore et al., 2012; Gogtay et al., 2004; Matsuzawa et al., 2001), our results suggest that frontal regions are involved in allocating attention in infancy.

Although we found recruitment of frontal regions in stimulus-driven attention, we observed noticeably weaker evidence in parietal regions, including the right TPJ. The TPJ allows the adult attention system to disengage from its current focus (Corbetta and Shulman, 2002), as would be needed to support reorientation in this task. One reason the TPJ may not have been involved more robustly is that TPJ is particularly important for goal-directed attention (de Fockert et al., 2004; Indovina and Macaluso, 2007; Kincade et al., 2005; Natale et al., 2010, 2009; Small et al., 2005), rather than stimulus-driven attention as tested here. However, infants have diminished goal-directed attention relative to adults (Csibra et al., 1998; Hood et al., 1998; Jakobsen et al., 2013; Scaife and Bruner, 1975). From this perspective, our results might offer new circumstantial evidence of the link between TPJ and goal-directed attention (Ellis and Turk-Browne, 2018). An alternative possibility is that the TPJ ROI was small and so may have been more affected by misalignment between participants. However, our supplemental analyses with dilated and spherical ROIs provide evidence against this possibility. Future studies could compare goal-directed and stimulus-driven attention using fMRI within the same infant participants to determine how these different modes of attention interact and develop.

In sum, we found that frontal regions from adult frontoparietal and cingulo-opercular networks are recruited to support stimulus-driven attention in infants. This study adds to the growing evidence that the frontal cortex supports infant cognition, despite undergoing substantial and protracted anatomical development (Ellis et al., 2020a; Raz and Saxe, 2020). Functionality of frontal cortex is honed over the course of development to support complex operations (Munakata et al., 2012), but these regions may be sufficiently developed in infancy to support stimulus-driven attention. This could reflect the importance of attention as a building block for learning and cognition, both in infancy and beyond.

## Methods

### Participants

Data from 24 sessions with infants aged 3.6 to 11.3 months (*M*=7.2; 14 female) met our minimum criterion for inclusion of four trials per condition. This sample does not include data from 11 sessions with participants in this age range that failed to produce enough data to reach criterion. Of the excluded sessions, three failed to meet the minimum number of trials and the rest collected enough trials but fell below criterion after exclusions for head motion and eye-tracking. In the final sample, four infants provided two sessions of usable data. These sessions typically occurred more than a month apart (range=0.9–3.0) and so the data were treated separately, similar to prior work (Deen et al., 2017). Of the 24 sessions, five were collected at the Magnetic Resonance Research Center (MRRC), and the rest were collected at the Brain Imaging Center (BIC). Refer to Table S1 for information on each participant. Parents provided informed consent on behalf of their child. The study was approved by the Human Investigation Committee and the Human Subjects Committee at Yale University.

### Materials

The code for running the cuing task can be found at: https://github.com/ntblab/experiment_menu. The code for the general analysis pipeline can be found here: https://github.com/ntblab/infant_neuropipe. The code for performing the specific analyses described in this paper can be found here: https://github.com/ntblab/infant_neuropipe/tree/PosnerCuing/. The data, including anonymized anatomical images, and both raw and preprocessed functional images can be found at: (to be shared on Dryad when published).

### Data acquisition

Data were acquired with a Siemens Prisma (3T) MRI at both the MRRC and BIC sites with the 20-channel Siemens head coil. Anatomical images were acquired with a T1-weighted PETRA sequence (TR_1_=3.32ms, TR_2_=2250ms, TE=0.07ms, flip angle=6°, matrix=320×320, slices=320, resolution=0.94mm iso, radial slices=30000). Functional images were acquired with a whole-brain T2* gradient-echo EPI sequence (MRRC: TR=2s, TE=28ms, flip angle=71°, matrix=64×64, slices=36, resolution=3mm iso, interleaved slice acquisition; BIC: identical except TE=30ms, slices=34).

### Procedures

Conducting fMRI research with awake infants is challenging for multiple reasons. Our protocol is described and validated in a separate methods paper (Ellis et al., 2020b). In brief, families visited the lab prior to their initial scanning session for an orientation session. This acclimated the infant and parent to the scanning environment. Scanning sessions were scheduled for a time when the parents felt the infant would be compliant. The infant and parent were extensively screened for metal. Hearing protection was applied to the infant in three layers: silicon inner ear putty, over-ear adhesive covers, and ear muffs. The infant was placed on the scanner bed, on top of a vacuum pillow that reduced movement comfortably. The top of the head coil was not used because the bottom elements provided sufficient coverage of the infant’s head. This increased visibility for monitoring infant comfort and allowed us to project stimuli onto the ceiling of the bore directly above the infant’s face using a custom mirror system. Using only the bottom of the head coil could have resulted in decreased data quality. However, detailed analyses revealed high-quality signal with this setup even in anterior regions of the infant brain farthest from the bottom head coil, likely as a result of their smaller head size (Ellis et al., 2020b). A video camera (MRRC: MRC 12M-i camera; BIC: MRC high-resolution camera) recorded the infant’s face during scanning for monitoring and eye-tracking.

When the infant was calm and focused, stimuli were shown in MATLAB using Psychtoolbox (http://psychtoolbox.org). The stimuli were pastel colored shapes (Johnson et al., 1994). Specifically, the attention-getter was a multi-colored square that oscillated in size from 2.5° to 7.5° at 1Hz. The cue was a green triangle that was 3° wide by 1.5° high. The target was a multi-colored pinwheel that rotated at 1Hz and subtended 10°.

Each trial began with the attention-getter. An experimenter in the control room monitored the participant’s eyes and when the participant was looking at the screen, the experimenter pressed a key to trigger a trial that began at the start of the next TR pulse. This meant the fixation was of variable length. Once triggered, the fixation was removed and the cue was shown for 100ms. The cue was presented 10° to the left, right, or both sides of fixation. After the cue offset, the screen was blank for 100ms before the target appeared. The target was also centered 10° either to the left or right of fixation. The target was on screen for 2s, after which the screen went blank until the next trial was initiated.

There were three trial types: valid, invalid and neutral. On valid trials, the cue and target were presented on the same side. On invalid trials, cue and target were presented on opposite sides. On neutral trials, cues were presented bilaterally and the target appeared at one of those locations. Trials were divided into blocks, each containing eight trials randomly intermixed (4 valid, 2 invalid, and 2 neutral), with cue and target side counterbalanced within each condition. Our goal was to collect four blocks per participant, but we collected more if we thought that a block might be unusable (*M*=4.7 blocks, range=4–7). There was at least 6s of rest between blocks where the screen was blank.

### Gaze coding

The gaze behavior of each infant was coded offline by two or three coders (*M*=2.3) who were blind to the condition. The coders determined whether the gaze was oriented ‘left’, ‘right’, ‘center’, ‘off-screen’ (i.e., blinking or looking away), or ‘undetected’ (i.e., out of the camera’s field of view or obscured by a hand or other object). For frames in which only the attention-getter was on the screen and nothing else, the coder was told that the infant was “probably looking at center”. This helped to calibrate the coder but did not prevent them deviating from the instruction if they were confident the child was looking elsewhere. Coders were instructed to label frames according to where they thought the eyes were directed. This protocol meant that coders often changed the label mid-saccade when the participant had looked left or right ‘enough’. We used this protocol to be consistent across experiments where the instantaneous position of the eye, rather than the trajectory, was important. We could have alternatively instructed coders to only change the label when the saccade was completed. Our protocol, compared to this alternative, would have shorter estimates of RT. Critically, this possibility does not introduce a bias between trial conditions.

Every video frame of each infant was coded at least once across coders. The frame rate and resolution varied by camera and site, but the minimum rate was 16Hz and the resolution was always sufficient to identify the eye. The label for each frame was determined as the mode of a moving window of five frames centered on that frame across all coder reports. In case of a tie, the modal response from the previous frame was used. The coders were highly reliable: When coding the same frame, coders reported the same response on 84% (range across participants=64–99%) of frames. Trials were excluded if the participant was not looking for the majority of the time during the cue presentation, if they looked to the target location before its onset, or if they did not look at the target within 1000ms of its onset.

### Preprocessing

Individual runs were preprocessed using FEAT in FSL (https://fsl.fmrib.ox.ac.uk/fsl), with a modified pipeline for infant data. Three volumes were discarded from the beginning of each run, in addition to the volumes automatically discarded by the EPI sequence. Blocks were stripped of any excess burn-in or burn-out volumes greater than the 3 TRs (6s) of rest after each block. Pseudo-runs were created if other experiments, not discussed here, were started in a run with the data of interest (sessions with a pseudo-run, N=20). Blocks were sometimes separated by long breaks (>30s) within a session because the participant was taken out of the scanner, because an anatomical scan was collected, or because of intervening experiments (N=8; *M*=391.1s break; range=63.7–1342.8s). The reference volume for alignment and motion correction was the ‘centroid’ volume that had the minimal Euclidean distance from all other volumes. The slices in each volume were realigned using slice-time correction. Time-points were excluded when there was greater than 3mm of movement from the previous time-point (*M*=11.7%, range=0.0–32.6%). We interpolated rather than removed these time-points so that they did not bias the linear detrending (in later analyses these time-points were ignored). Blocks were excluded if more than 50% of the time-points were excluded. To determine which voxels were brain and which were non-brain we created a mask from the signal-to-fluctuating-noise ratio (SFNR) for each voxel in the centroid volume. The data were smoothed with a Gaussian kernel (5mm FWHM) and linearly filtered in time. AFNI’s (https://afni.nimh.nih.gov) despiking algorithm was used to attenuate aberrant time-points within voxels. For further explanation and justification of this preprocessing procedure, refer to (Ellis et al., 2020b).

We aligned each run’s centroid volume to the infant’s anatomical scan from the same session. We used FLIRT with a normalized mutual information cost function to create the initial alignment. Additional manual alignment was then performed using mrAlign from mrTools (Gardner lab) to fix deficiencies of automatic alignment. The preprocessed functional data were aligned into anatomical space but retained their original spatial resolution (3mm iso). The anatomical scan from each participant was automatically (FLIRT) and manually (Freeview) aligned to an age-specific MNI infant template (Fonov et al., 2011) and then aligned to the adult MNI template (MNI152). This allowed the functional data to be transformed into standard space. ROI and whole-brain voxelwise analyses were performed in this 1mm MNI space. To determine which voxels to consider at the group level, the intersection of brain voxels from all infant participants in standard space was used as a mask.

Participants were considered usable if they had at least four usable trials of each condition. In the final sample, and consistent with the frequency of trial types, participants had an average of 12.9 (range: 8–18) valid trials, 6.7 (range: 4–10) neutral trials, and 6.6 (range: 4–10) invalid trials.

To account for differences in intensity and variance across runs, the blocks that survived exclusions were normalized over time within run using *z*-scoring, prior to the runs being concatenated for further analyses.

### Behavior analysis

We quantified the RT for the participant to saccade to the target. In particular, the onset time of the target was subtracted from the time-stamp of the frame when the participant first looked to the correct side (i.e., left if the target was on the left). Only trials that met the inclusion criteria (e.g., low head motion, looking during the cue presentation) were retained for analysis. The RT for each trial was averaged within condition and then compared across participants. Non-parametric bootstrap resampling was used to compare the conditions (Efron and Tibshirani, 1986). Namely, for each test we sampled 24 participants with replacement 10,000 times, computing the mean across participants on each iteration to generate a sampling distribution. For null hypothesis testing, we calculated the *p*-value as the proportion of samples where the mean was in the opposite direction to the true effect, doubled to make the test two-tailed. A similar bootstrap resampling procedure was used to statistically evaluate the correlation between age and behavior: the age and behavior bivariate data from each participant was sampled with replacement 10,000 times, and for each sample the Pearson correlation was calculated. The *p*-value was the proportion of samples resulting in a correlation with the opposite sign from the true correlation, doubled to make the test two-tailed. We also performed a follow-up analysis using only the first block of usable data. One participant did not have any usable invalid trials in their first block so they were not included in statistical comparisons involving invalid trials in this analysis.

### GLM analysis

For the main analysis, a GLM was fit to the preprocessed and *z*-scored BOLD activity using FEAT in FSL. Separate regressors were specified for valid, invalid and neutral trials. An event of a given condition began at the onset of the cue and ended at the target offset. Each event was modeled as a boxcar (2.2s duration), convolved with a double-gamma hemodynamic response function. The six translation and rotation parameters from motion correction were included in the GLM as regressors of no interest. Excluded TRs were scrubbed with an additional regressor for each to-be-excluded time-point (Siegel et al., 2014). The main contrasts compared invalid greater than valid trials, neutral greater than valid trials, and invalid greater than neutral trials. The *z*-statistic volumes corresponding to these three contrasts were extracted for each participant and aligned to standard space where subsequent analyses were performed. To visualize each contrast, we performed a voxelwise whole-brain *t*-test across participants.

ROIs were defined with Neurosynth, a meta-analytic tool for identifying loci of activation from published fMRI studies (Yarkoni et al., 2011). Specifically, we acquired the statistic map for the term “attention” on December 1, 2019. This aggregated across 1831 studies and used a False Discovery Rate of 0.01 to detect regions that were reliably implicated in attention. Using FSL’s cluster algorithm, we identified 9 clusters with at least 27 voxels: in ascending size, rTPJ, lIPS, rLOC, lLOC, rACC, lFEF, lSPL, rFEF, and rSPL (Table S2). Lower-case ‘r’ and ‘l’ indicate left and right hemisphere, respectively. Because this is a functional atlas, rather than an anatomical atlas, some of these regions do not map cleanly onto anatomy but the names were chosen to be consistent with other regions described in the literature. For instance, the ACC ROI includes large portions of the paracingulate cortex and may be instead described as medial cingulate cortex, which is part of the dorsal salience network (Touroutoglou et al., 2012). The atlas was in 2mm isotropic space and so we upsampled it to 1mm space to be consistent with our other data. We manually divided two of the ROIs since they included anatomically and theoretically distinct regions. Namely, we split off part of rFEF to make rMFG and part of rSPL to make rIPS. We collapsed ROIs bilaterally when available, such as lFEF and rFEF. The mean *z*-statistic value across voxels within each ROI was computed for each participant and contrast. The statistical significance of each ROI and correlations with age were evaluated using bootstrap resampling (as described above). For follow-up analyses, we edited these ROIs in two ways. First, we performed one step of modal dilation to increase the size of each region while preserving its overall shape. Second, we created spherical ROIs around the peak voxel within each ROI. These spheres had a radius of 10mm, resulting in a constant voxel count across regions, although the shape did not reflect the original ROI.

### Leave-one-participant-out ROI analysis

To identify clusters in a data-driven manner, we used a leave-one-participant-out approach (Esterman et al., 2010). This allowed us to identify regions reliably recruited in infants while making as few assumptions as possible about where the activity will localized from adults. Because one participant is held-out, each participant can be treated as independent, akin to cross-validation (Kriegeskorte et al., 2009). As an illustrative example from this dataset, the right ACC was identified in 12/24 leave-one-participant-out iterations for the invalid greater than valid contrast. Yet, an example voxel (MNI: −4, 36, 16) in this cluster showed greater activity in invalid than valid trials in 18/24 participants. Indeed, of the 12 participants for whom this region was not identified (i.e., when they were left out, the cluster from other participants did not form in this region), eight showed greater activity in invalid than valid trials in this same voxel. Hence, this is a conservative, data-driven method that requires consistency within the group.

To implement this approach, the *z*-statistic volumes (e.g., invalid greater than valid) from running a GLM on all but one participant were concatenated and a *t*-test was performed. Voxels that had a *t*-statistic exceeding a two-tailed *p*-value of 0.01 were then clustered such that only clusters with at least 27 voxels survived. These criteria were chosen to produce sufficiently small clusters and to be consistent with how the functional ROIs were constructed. The voxel with the peak *t*-statistic in each cluster was then labeled according to the Harvard-Oxford cortical atlas (Makris et al., 2006). If this voxel was in a cortical region with at least 50% probability then this label was used, if not, then a subcortical label was used. Clusters labelled in a ventricle were ignored.

The effect for each cluster was then quantified in the held-out participant by averaging their contrast values across the voxels in the cluster. These results were pooled and compared across participants if the cluster had the same anatomical label (as the clusters on each iteration were in slightly different locations). To be considered for statistical analysis, clusters with the same label had to be found in at least half of the participants. We then performed bootstrap resampling with each label that exceeded that threshold. Because this analysis was exploratory, we used a Holms-Bonferroni correction on the alpha value.

### Saccade-evoked response

We examined whether the saccadic behavior differed between conditions and whether this could explain our results. We first quantified every change in coded location between the ‘left’, ‘center’, and ‘right’(changes in gaze involving ‘off-screen’ or ‘undetected’ were ignored). The average number of saccades per condition was compared using bootstrap resampling. We then created a boxcar out of these saccades, with each saccade having a duration of 100ms, and convolved the boxcar with a double-gamma hemodynamic response to create a regressor. We used that regressor to predict fMRI activity in the GLM and extracted the whole-brain *z*-statistic map. The *z*-statistics were averaged across voxels in each ROI.

## Acknowledgements

We thank all of the families who participated; N. Wilson, N. Córdova and V. Bejjanki for help with initial set up; K. Armstrong, C. Greenberg, J. Bu, L. Rait, J. Daniels, and the entire Yale Baby School team for recruitment, scheduling, and administration; H. Faulkner, Y. Braverman, J. Fel, and J. Wu for help with gaze coding; and R. Lee, L. Nystrom, N. DePinto, and R. Watts for technical support. We are grateful for internal funding from the Department of Psychology and Princeton Neuroscience Institute at Princeton University and from the Department of Psychology and Faculty of Arts and Sciences at Yale University. N.B.T-B. was further supported by the Canadian Institute for Advanced Research.

---

**Figure S1:**
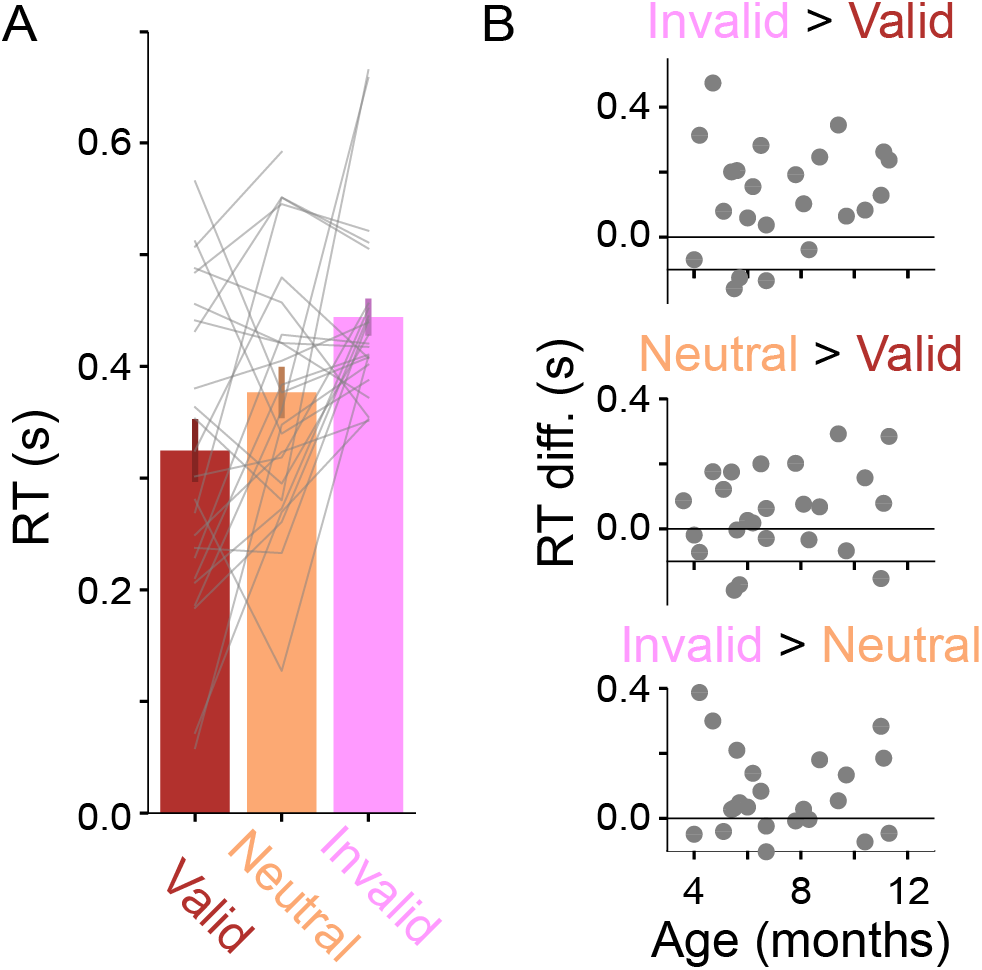
Analysis of attention behavior in the first usable block from each participant. Note that one participant did not have any usable invalid trials in their first block so they were excluded from comparisons involving invalid trials. (A) Average RT in seconds for each trial type. Invalid trials were slower than valid trials (invalid>valid in 18/23 participants, *M*=0.13s, CI=[0.06, 0.19], *p>*0.001), neutral trials were slower than valid trials (neutral>valid in 15/24 participants, *M*=0.05s, CI=[0.00, 0.10], *p*=0.049), and neutral trials were faster than invalid trials (invalid>neutral in 15/23 participants, *M*=0.08s, CI=[0.03, 0.13], *p*=0.002). Error bars indicate standard error across participants. Gray lines connect individual participant data across the three trial types. (B) The differences between conditions were not reliably related to participant age (invalid<valid: r=0.15, *p*=0.479; neutral<valid: r=0.21, *p*=0.352; invalid<invalid: r=−0.07, *p*=0.804).

**Figure S2:**
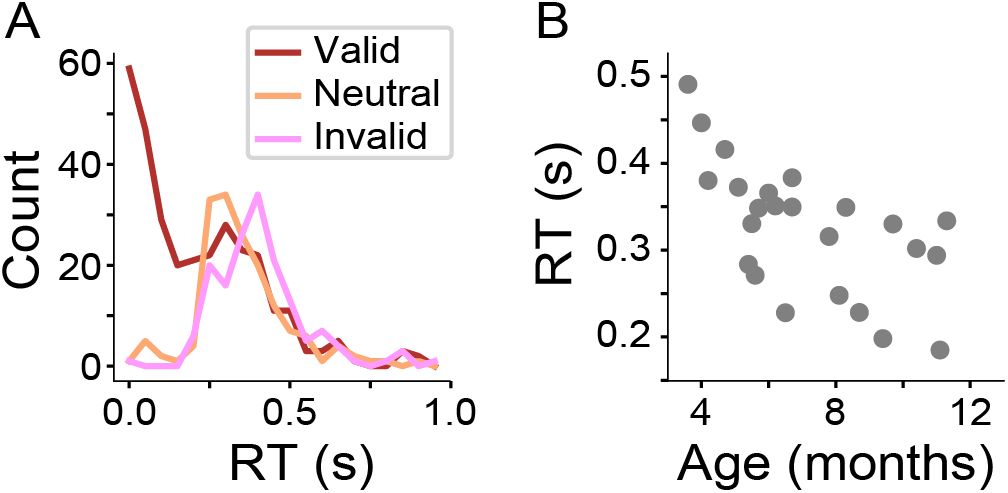
Information about RT behavior. (A) Because we could not instruct the infants to maintain fixation until the target appeared, it was possible for them to initiate the saccade as soon as the cue appeared. To explore this, we pooled all trials from all participants and created a histogram of RTs for each condition. (B) We also examined how the average RT across trial types for each participant related to age. As participants got older, they were faster to saccade to the target (r=−0.63, *p*<0.001).

**Figure S3:**
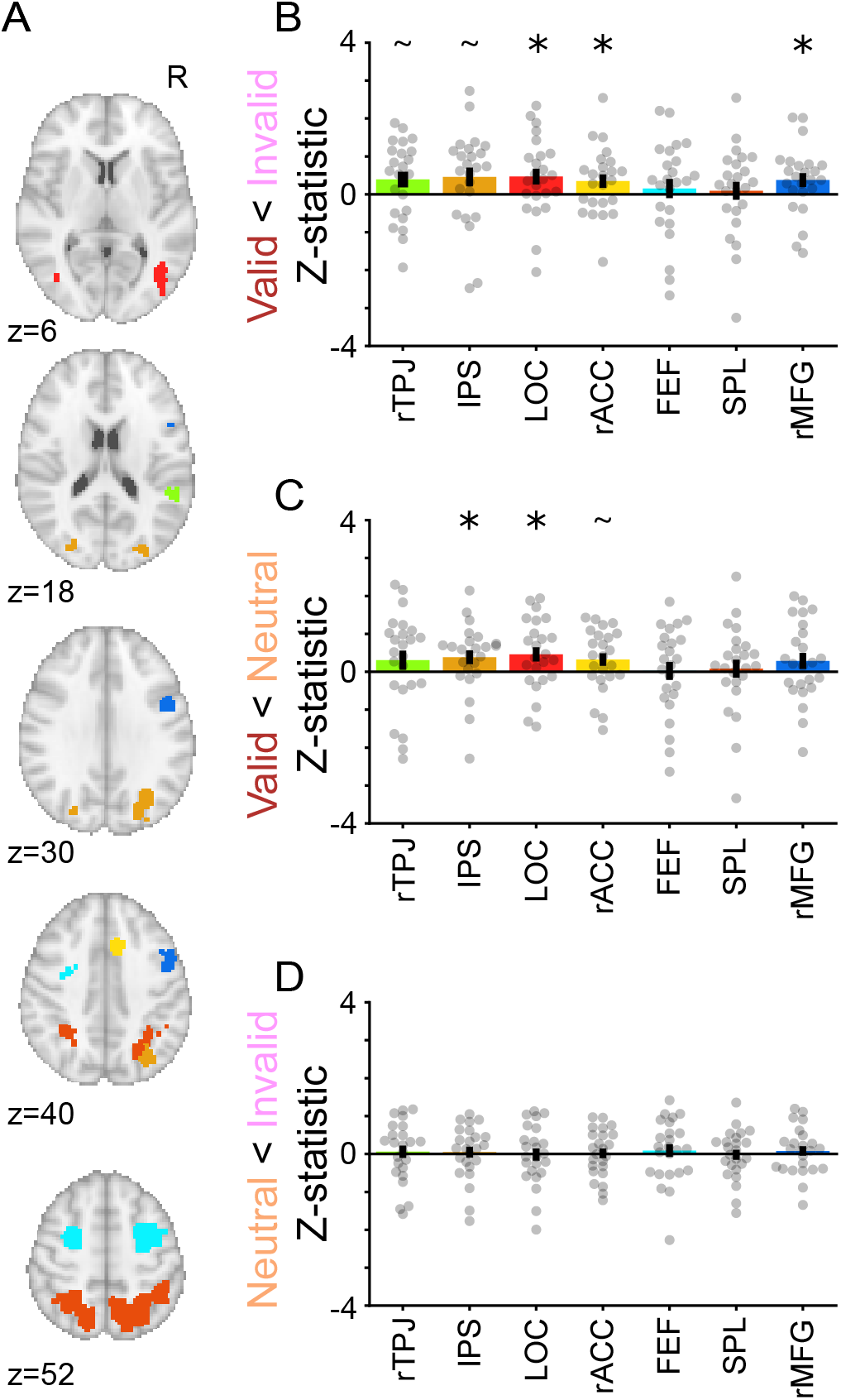
Re-analysis with dilated ROIs. (A) ROIs from functional atlas (Figure 3A) but increased in size by one step of modal dilation (on average 1 voxel on every edge). (B) The contrast of valid and invalid trials resulted in significantly higher activity for invalid trials in LOC (*M*=0.47, CI=[0.06, 0.87], *p*=0.029), rACC (*M*=0.35, CI=[0.02, 0.69], *p*=0.040), and MFG (*M*=0.37, CI=[0.02, 0.73], *p*=0.040), and marginally higher activity in rTPJ (*M*=0.39, CI=[−0.03, 0.78], *p*=0.069) and IPS (*M*=0.46, CI=[-0.05, 0.93], *p*=0.073). (C) The contrast of valid and neutral trials resulted in significantly higher activity for neutral trials in IPS (*M*=0.38, CI=[0.01, 0.72], *p*=0.043) and LOC (*M*=0.46, CI=[0.08, 0.83], *p*=0.018), and marginally higher activity in rACC (*M*=0.32, CI=[−0.02, 0.64], *p*=0.062). (D) The contrast of invalid and neutral trials resulted in no significant regions. Lower-case ‘r’ and ‘l’ indicate left and right hemisphere, respectively. Error bars indicate standard error across participants. *=*p*<0.05, ^~^=*p*<0.075

**Figure S4:**
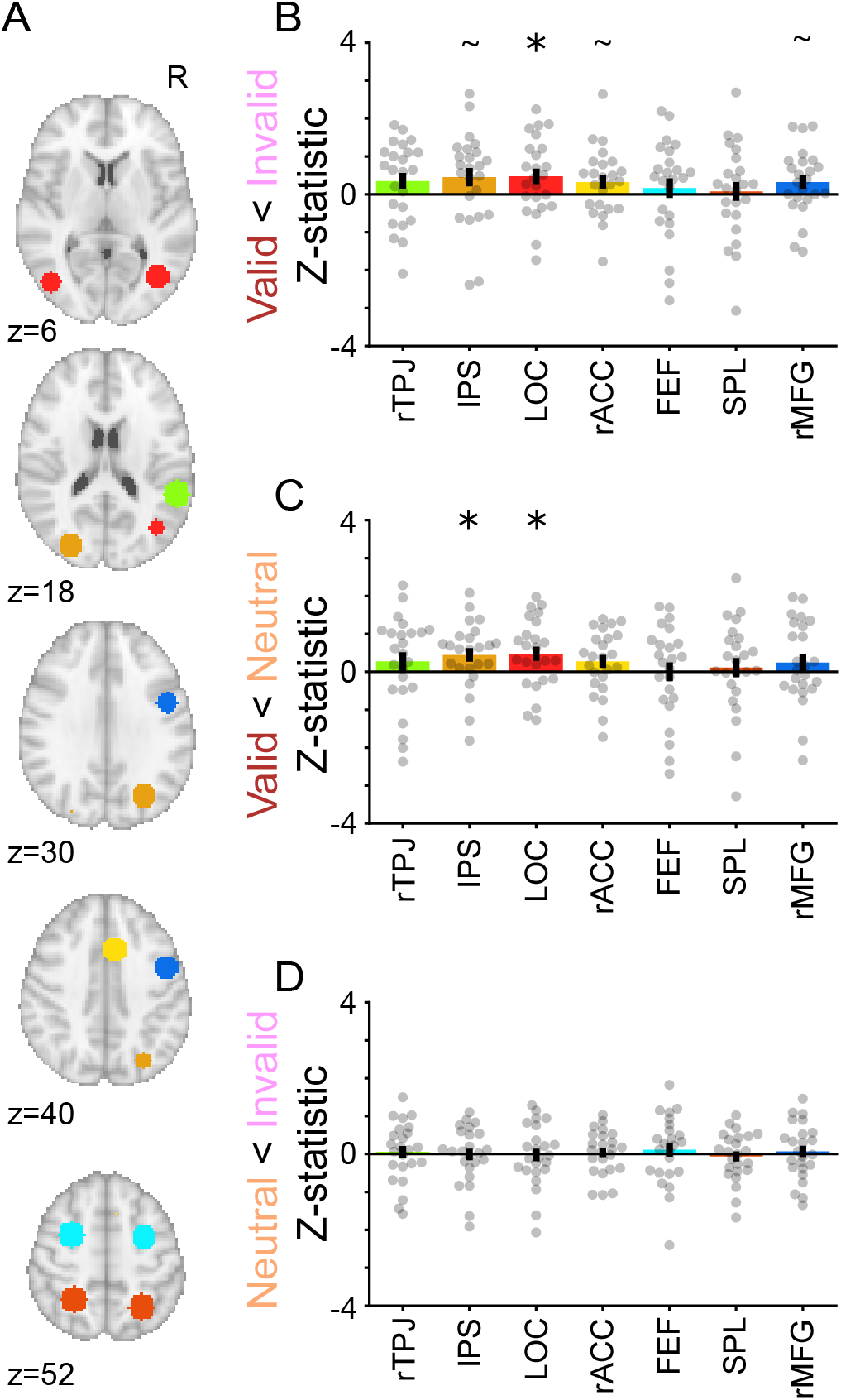
Re-analysis with spherical ROIs. (A) ROIs from functional atlas (Figure 3A) but with a sphere of radius 10mm drawn around the peak voxel. (B) The contrast of valid and invalid trials resulted in significantly higher activity for invalid trials in LOC (*M*=0.48, CI=[0.07, 0.87], *p*=0.020), and marginally higher activity in IPS (*M*=0.45, CI=[−0.04, 0.91], *p*=0.069), rACC (*M*=0.32, CI=[−0.02, 0.68], *p*=0.060), and rMFG (*M*=0.32, CI=[−0.03, 0.67], *p*=0.070). (C) The contrast of valid and neutral trials resulted in significantly higher activity for neutral trials in IPS (*M*=0.45, CI=[0.09, 0.78], *p*=0.013), and LOC (*M*=0.48, CI=[0.11, 0.84], *p*=0.012). (D) The contrast of invalid and neutral trials resulted in no significant regions. Lower-case ‘r’ and ‘l’ indicate left and right hemisphere, respectively. Error bars indicate standard error across participants. *=*p*<0.05, ^~^=*p*<0.075

**Figure S5:**
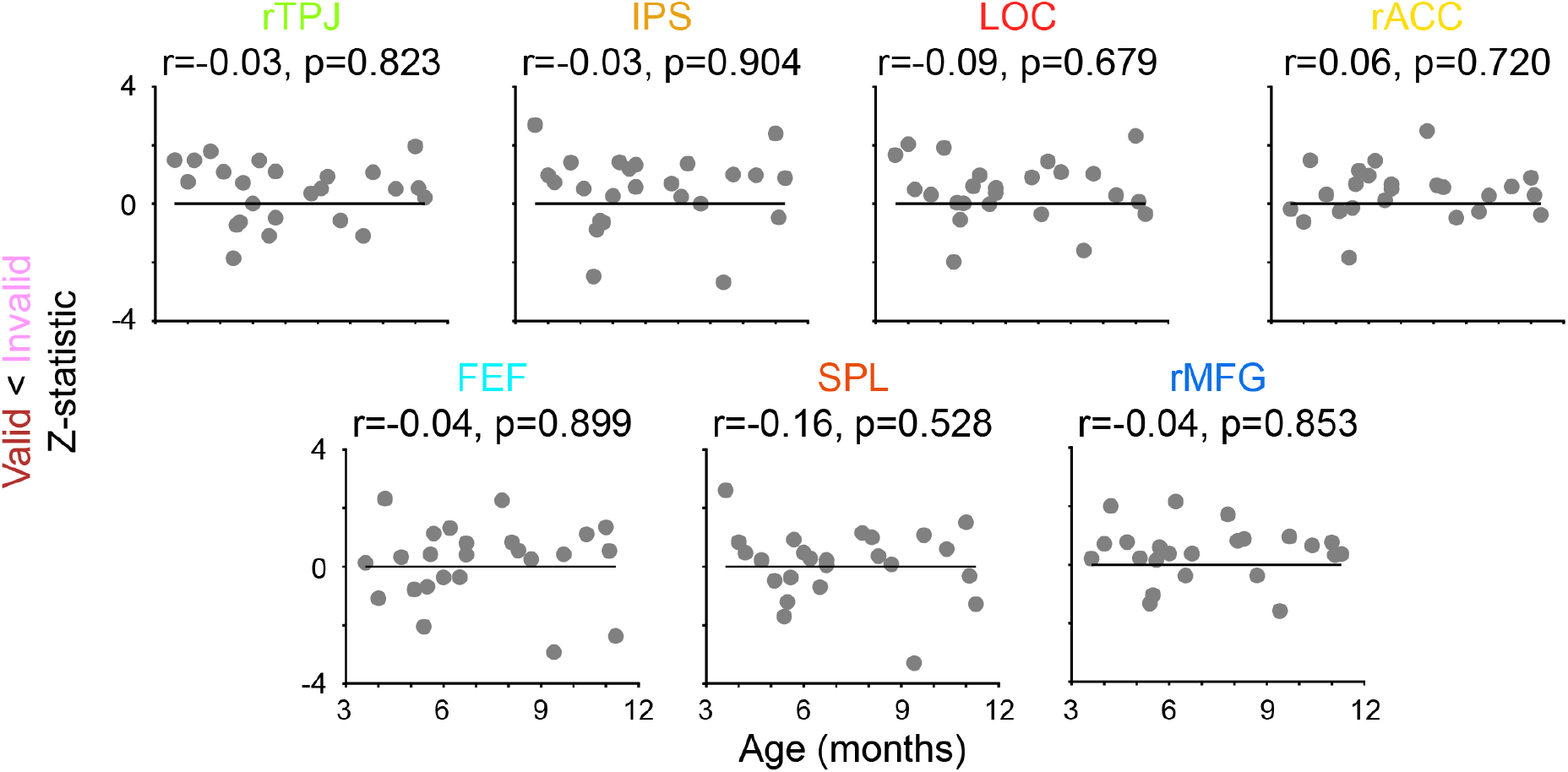
Relationship between participant age and the invalid>valid contrast from Figure 3A.

**Figure S6:**
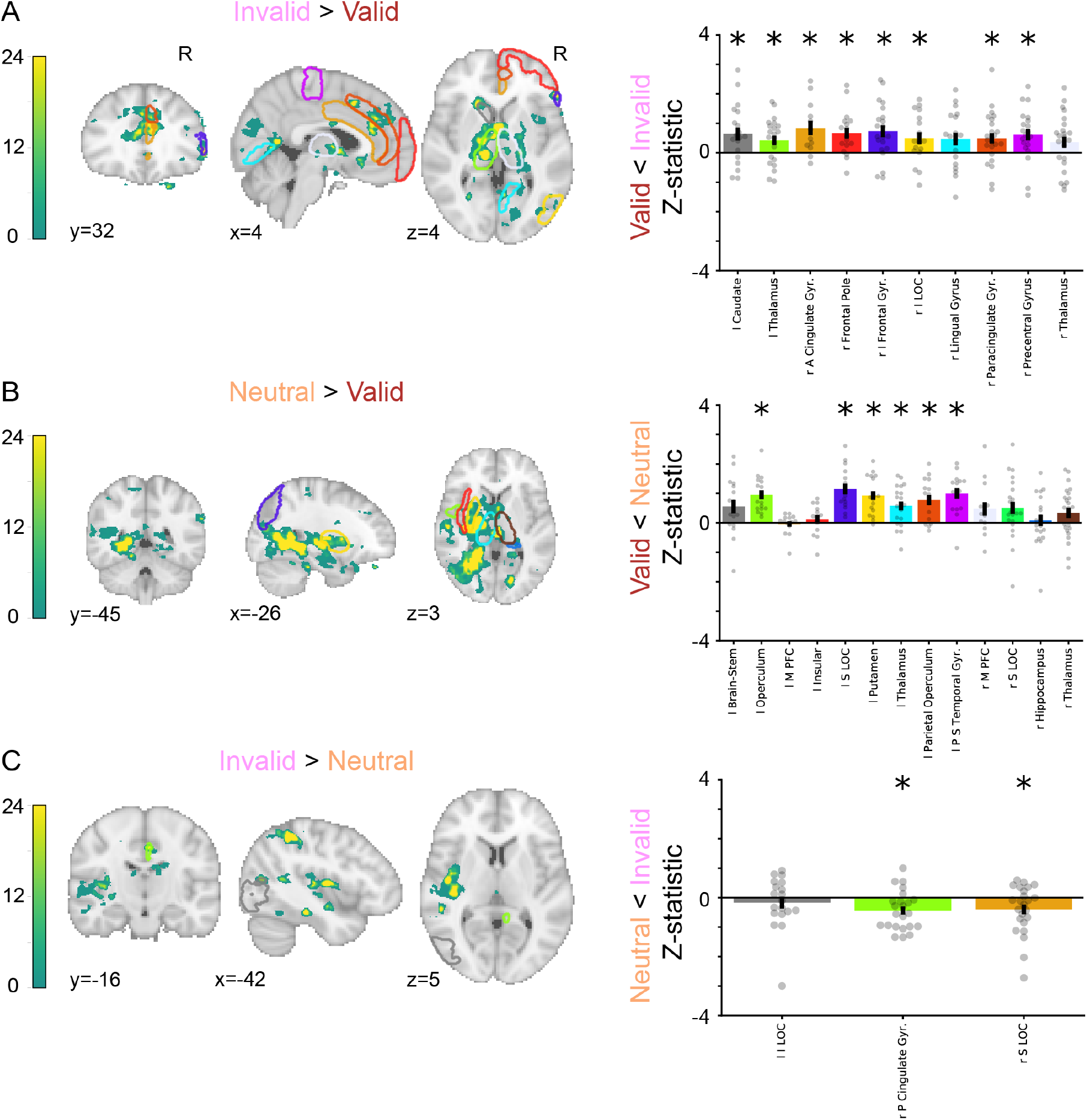
Leave-one-participant-out ROI analyses. The three trial type contrasts are shown: (A) invalid>valid, (B) neutral>valid, and (C) invalid>neutral. Left panels show the count of voxels that form clusters on each leave-one-participant-out fold. Overlaid on these figures are the ROIs (in MNI space) that are found in at least half of the training sets. Right panels show the mean activity across voxels in each cluster from the held-out participant. For the ROI names: ‘l’=left hemisphere, ‘r’=right hemisphere, ‘A’=anterior, ‘P’=posterior, ‘I’=inferior, ‘S’=superior, ‘Gyr’=gyrus. *=*p*<0.05 after Holms-Bonferroni

**Figure S7:**
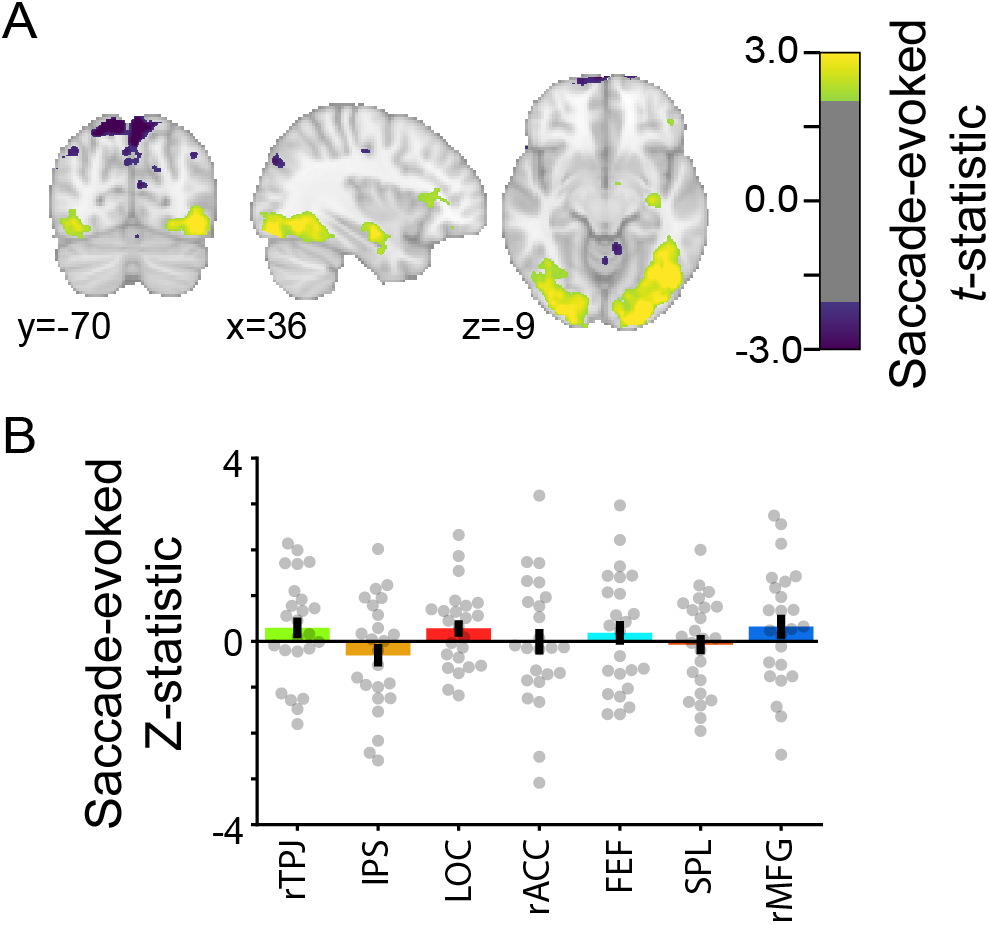
Saccade-evoked activity during the task. A regressor was created such that any saccades that occurred during a trial were treated as an event lasting 100ms, and these events were convolved with a double-gamma hemodynamic response function. (A) Saccades evoked widespread activity in early visual cortex (*p*<0.05 uncorrected; coordinates in MNI space). (B) However, saccades did not produce reliable responses in the main ROIs (Figure 3A), several of which distinguished attention conditions.

**Table 1:** Demographic information. ‘ID’ is a unique infant identifier (i.e., sXXXX Y Z), with the first four digits (XXXX) indicating the family, the fifth digit (Y) the child number within family, and the sixth digit (Z) the session number with that child. ‘Age’ is recorded in months. ‘Sex’ is female or male. ‘Site’ is Magnetic Resonance Research Center (MRRC) or Brain Imaging Center (BIC). ‘Blocks’ is the number of total blocks participant saw. Number of usable trials per condition are provided. ‘Prop. incl. TR’ is the proportion of TRs included from usable blocks. ‘Prop. incl. frames’ is the proportion of eye tracking data included from usable blocks. ‘Num. coders’ is the number of gaze coders per participant.

| ID | Age | Sex | Site | Blocks | Valid trials | Neutral trials | Invalid trials | Prop. incl. TRs | Prop. incl. frames | Num. coders |
| --- | --- | --- | --- | --- | --- | --- | --- | --- | --- | --- |
| s6057_1.1 | 3.6 | M | BIC | 4 | 9 | 7 | 6 | 0.86 | 0.72 | 2 |
| s6607_1.1 | 4.0 | M | MRRC | 4 | 15 | 7 | 5 | 0.98 | 0.79 | 2 |
| s7037_1.1 | 4.2 | M | BIC | 7 | 10 | 6 | 5 | 0.68 | 0.76 | 2 |
| s9057_1.1 | 4.7 | F | BIC | 4 | 8 | 4 | 6 | 0.82 | 0.86 | 2 |
| s1607_1.1 | 5.1 | M | MRRC | 5 | 10 | 6 | 7 | 0.68 | 0.64 | 3 |
| s8047_1.1 | 5.4 | F | BIC | 4 | 13 | 7 | 7 | 1.00 | 0.90 | 2 |
| s0057_1.1 | 5.5 | F | BIC | 6 | 17 | 10 | 7 | 0.93 | 0.71 | 3 |
| s2037_1.1 | 5.6 | M | BIC | 5 | 17 | 6 | 7 | 0.85 | 0.85 | 2 |
| s3687_1.2 | 5.7 | F | MRRC | 4 | 9 | 5 | 6 | 0.81 | 0.79 | 3 |
| s2687_1.2 | 6.0 | M | MRRC | 4 | 14 | 8 | 6 | 0.88 | 0.92 | 2 |
| s7067_1.2 | 6.2 | F | BIC | 4 | 13 | 8 | 8 | 1.00 | 0.94 | 2 |
| s2037_1.2 | 6.5 | M | BIC | 5 | 18 | 9 | 10 | 0.93 | 0.89 | 2 |
| s3017_1.1 | 6.7 | M | BIC | 4 | 11 | 5 | 6 | 0.98 | 0.90 | 2 |
| s4607_1.2 | 6.7 | F | MRRC | 4 | 14 | 5 | 7 | 0.98 | 0.92 | 3 |
| s0057_1.2 | 7.8 | F | BIC | 5 | 10 | 5 | 6 | 0.67 | 0.75 | 2 |
| s6067_1.1 | 8.1 | M | BIC | 5 | 9 | 6 | 6 | 0.70 | 0.93 | 2 |
| s7017_1.4 | 8.3 | F | BIC | 4 | 15 | 8 | 6 | 0.93 | 0.91 | 2 |
| s5037_1.1 | 8.7 | M | BIC | 5 | 16 | 8 | 10 | 0.99 | 0.89 | 2 |
| s1057_1.2 | 9.4 | F | BIC | 5 | 12 | 6 | 4 | 0.79 | 0.91 | 2 |
| s4607_1.3 | 9.7 | F | BIC | 5 | 10 | 6 | 5 | 0.92 | 0.75 | 2 |
| s3607_1.2 | 10.4 | F | BIC | 4 | 14 | 7 | 7 | 0.96 | 0.90 | 3 |
| s8687_1.3 | 11.0 | F | BIC | 4 | 13 | 6 | 6 | 0.97 | 0.90 | 3 |
| s1057_1.3 | 11.1 | F | BIC | 7 | 16 | 7 | 7 | 0.91 | 0.95 | 2 |
| s6687_1.3 | 11.3 | F | BIC | 5 | 17 | 9 | 9 | 0.97 | 0.73 | 3 |
| Av. | 7.15 | . | . | 4.71 | 12.92 | 6.71 | 6.62 | 0.88 | 0.84 | 2.29 |

**Table 2:** ROI information. MNI coordinates and region size for each ROI from the functional atlas. ROIs that are collapsed bilaterally for analysis (e.g., lFEF and rFEF) are listed separately here. Volume is provided in cubic millimeters.

| Name | X | Y | Z | Volume |
| --- | --- | --- | --- | --- |
| rTPJ | 54 | -42 | 18 | 256 |
| rIPS | 28 | -70 | 32 | 2526 |
| lIPS | -28 | -82 | 20 | 280 |
| rLOC | 38 | -68 | 10 | 664 |
| lLOC | -44 | -72 | 0 | 672 |
| rACC | 6 | 16 | 42 | 736 |
| rFEF | 28 | -2 | 54 | 3120 |
| lFEF | -28 | 0 | 52 | 1720 |
| rSPL | 26 | -56 | 52 | 8226 |
| lSPL | -26 | -50 | 52 | 4432 |
| rMFG | 46 | 2 | 36 | 1552 |

